# Single-cell sequencing reveals αβ chain pairing shapes the T cell repertoire

**DOI:** 10.1101/213462

**Authors:** Kristina Grigaityte, Jason A. Carter, Stephen J. Goldfless, Eric W. Jeffery, Ronald J. Hause, Yue Jiang, David Koppstein, Adrian W. Briggs, George M. Church, Francois Vigneault, Gurinder S. Atwal

## Abstract

A diverse T cell repertoire is a critical component of the adaptive immune system, providing protection against invading pathogens and neoplastic changes, relying on the recognition of foreign antigens and neoantigen peptides by T cell receptors (TCRs). However, the statistical properties and function of the T cell pool in an individual, under normal physiological conditions, are poorly understood. In this study, we report a comprehensive, quantitative characterization of the T cell repertoire from over 1.9 million cells, yielding over 200,000 high quality paired αβ sequences in 5 healthy human subjects. The dataset was obtained by leveraging recent biotechnology developments in deep RNA sequencing of lymphocytes *via* single-cell barcoding in emulsion. We report non-random associations and non-monogamous pairing between the α and β chains, lowering the theoretical diversity of the T cell repertoire, and increasing the frequency of public clones shared among individuals. T cell clone size distributions closely followed a power law, with markedly longer tails for CD8^+^ cytotoxic T cells than CD4^+^ helper T cells. Furthermore, clonality estimates based on paired chains from single T cells were lower than that from single chain data. Taken together, these results highlight the importance of sequencing αβ pairs to accurately quantify lymphocyte receptor diversity.

## INTRODUCTION

The adaptive immune system of jawed vertebrates requires maintenance of a genetically diverse T cell repertoire in order to defend against the wide spectrum of potential antigens, whether infectious or neoplastic, that an individual may encounter over the course of their life. However, the statistical properties of the αβ T cell repertoire in healthy individuals has remained poorly characterized, in large part due to the laborious task of sequencing single T cells in a high throughput fashion. The TCR is a heterodimer consisting of one α and one β chain, each of which is highly variable. The germline DNA sequence of human α and β chains resides on differing chromosomes and encodes a number of genes – 60-70 variable (V) and 61 joining (J) genes for the α chain, and 52 V, 13 J, and 2 diversity (D) genes for the β chain (Fig. 1A,B). The DNA of each differentiated T cell undergoes a V(D)J recombination in the thymus - one Vα, Jα, Vβ, Dβ, and Jβ gene are selected via DNA splicing followed by random repair, insertions and deletions at the splice sites. Due to this mechanism, most of the genetic variation of the TCR repertoire resides in the complementarity-determining region 3 (CDR3) encompassing the V(D)J junction. The diversity of the T cell repertoire is further increased by the pairing of α and β chains, potentially generating over 10^15^ distinct TCRs^1–3^. Thus, an understanding of the full TCR repertoire requires the identification of paired αβ sequences, which has remained technically challenging.

**FIGURE 1.**
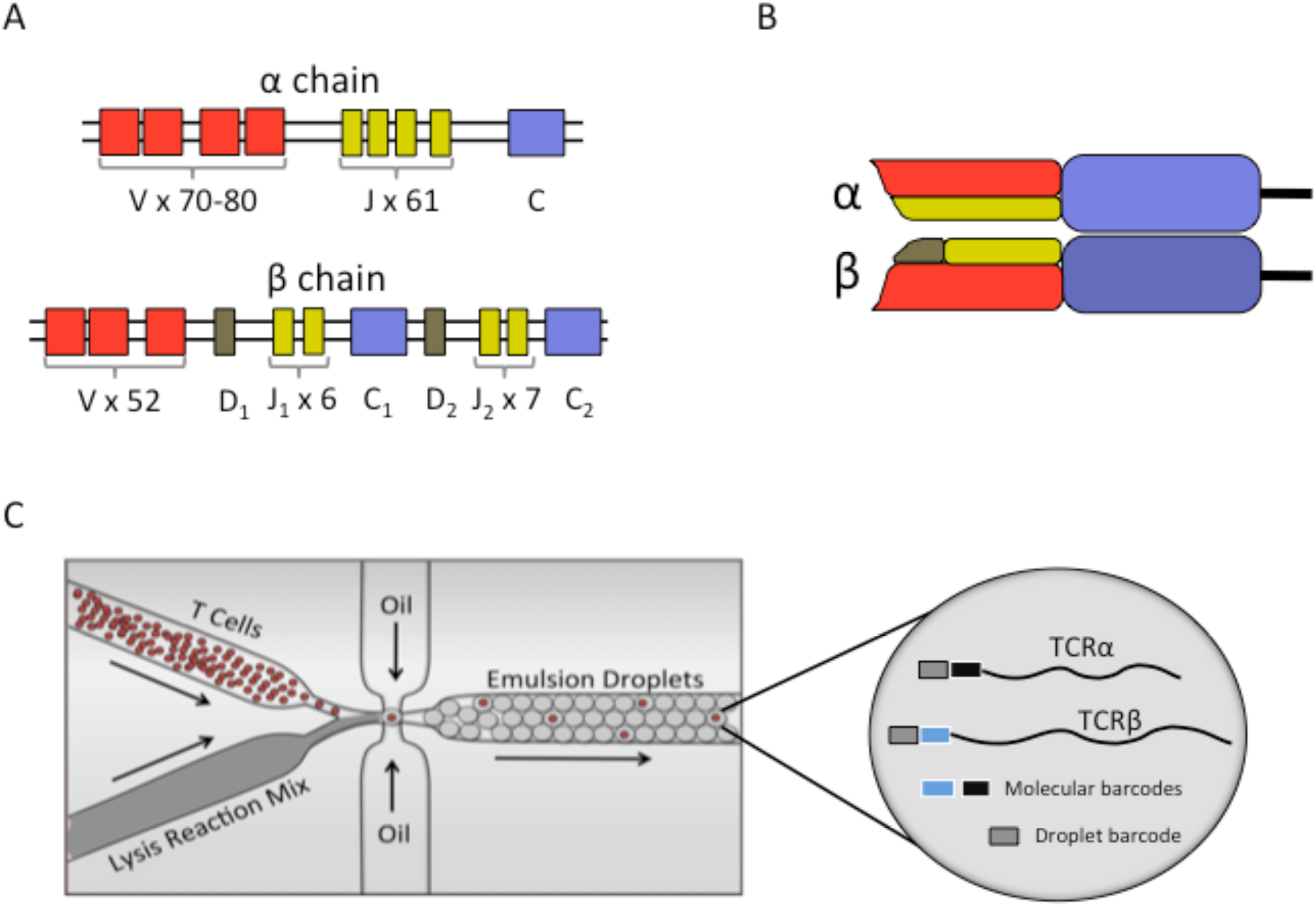
T cell receptors (TCRs) and high-throughput single-cell barcoding in emulsion. A) Human germline DNA depiction of TCR α and β chains. V – variable gene; J – junction gene; D – diversity gene; C – constant region. B) Functional T cell receptor structure. Colors correspond to germline gene notation in (A). C) High throughput single-cell barcoding in emulsion platform. Single T cells are encapsulated in oil droplets at an approximate Poisson rate of 0.1 cells/droplet to reduce the chance of multiple cells per droplet. This results in ~10% of droplets containing at least 1 T cell, and ~90% of cell containing droplets will have only one T cell. Droplet barcodes are attached to the 5’end of α and β sequences, allowing the recovery of the pairs post-sequencing.

Most studies of the T cell repertoire have, to date, focused on characterizing either the α or β chain diversity alone by bulk sequencing methods^4–9^. While convenient and cheaper, subsequent clonality analyses of bulk sequencing data necessarily invoke the commonly made assumption that the β chain is independent of the concomitant α chain^10–14^, despite a lack of direct evidence supporting this claim. Prior experimental attempts to recover αβ pairs involve single-cell sequencing in 96-well plates^15^, RT-PCR^16–19^ or sequencing in emulsion^20^, resulting in relatively low sample sizes and loss of rare T cell clones. Computational methods such as PairSEQ^21^ and ALPHABETR^22^ can overcome small sample sizes and have a reported capacity of recovering up to 10^5^ αβ pairs. Nevertheless, due to the probabilistic nature of these statistical methods, detecting small size clones still remains problematic, and an accurate description of the T cell repertoire within and across individuals has remained elusive.

Here, we analyzed an unprecedented high-throughput dataset of full-length, high quality, paired αβ sequences (n=205,950) from peripheral blood samples of 5 healthy individuals (3 males, 2 females, ages 33-69) acquired through a recently developed microfluidic method of single-cell RNA sequencing in emulsion droplets^23^. In brief, αβ pairs were recovered by attaching unique droplet and molecular barcodes to the target cDNA from the lysed cells inside droplets (Fig 1C), followed by recovery and next generation sequencing. The sequenced T cells were further stratified into CD4^+^ (n=73,495) and CD8^+^ (n=30,321) subtypes (Sup Table 1), based on paired sequence tags introduced by labeling with DNA-conjugated antibodies. All TCR sequences used in this study came from T cells expressing only one unique TCR. That is, any cell expressing more than one α or β chain, whether biological due to violations of allelic exclusion^24^ or artifactual due to multiple cells randomly included in a single droplet, were removed from the final dataset (see Methods).

## RESULTS

### Single chain α or β repertoires are not equivalent to the paired αβ repertoire

First, we address the assumption that every αβ pair is unique, i.e. that no *α* will pair with more than one different β and vice versa^10,11^. To evaluate potential discrepancies between paired and single chain analyses, we split our paired αβ-CDR3 dataset into separate α-CDR3 and β-CDR3 datasets, and compared clone size distributions of all three groups. That is, if we observe an αβ-CDR3 with a clone size 50, we create corresponding α-CDR3 and β-CDR3 clones of size 50 each. It is important to note here that each sample is of the same size, and therefore all differences observed arise solely from non-random αβ pairings rather than as an artifact of varying sample sizes. We find that the clone size frequency distributions of both single and paired αβ chains resemble heavily tailed power law distributions characterized by linear behavior on the bilogarithmic scale (see Methods; Eq 1). Given the unique αβ pairing assumption, one would expect such analyses to result in three identical distributions. However, we observe three distinct clone size distributions of paired αβ-CDR3, single chain α-CDR3, and single chain β-CDR3. This contradicts the assumption of unique pairing and therefore is suggestive of a non-random process driving αβ pairing (Figure 2A, Sup Fig. 1).

**FIGURE 2.**
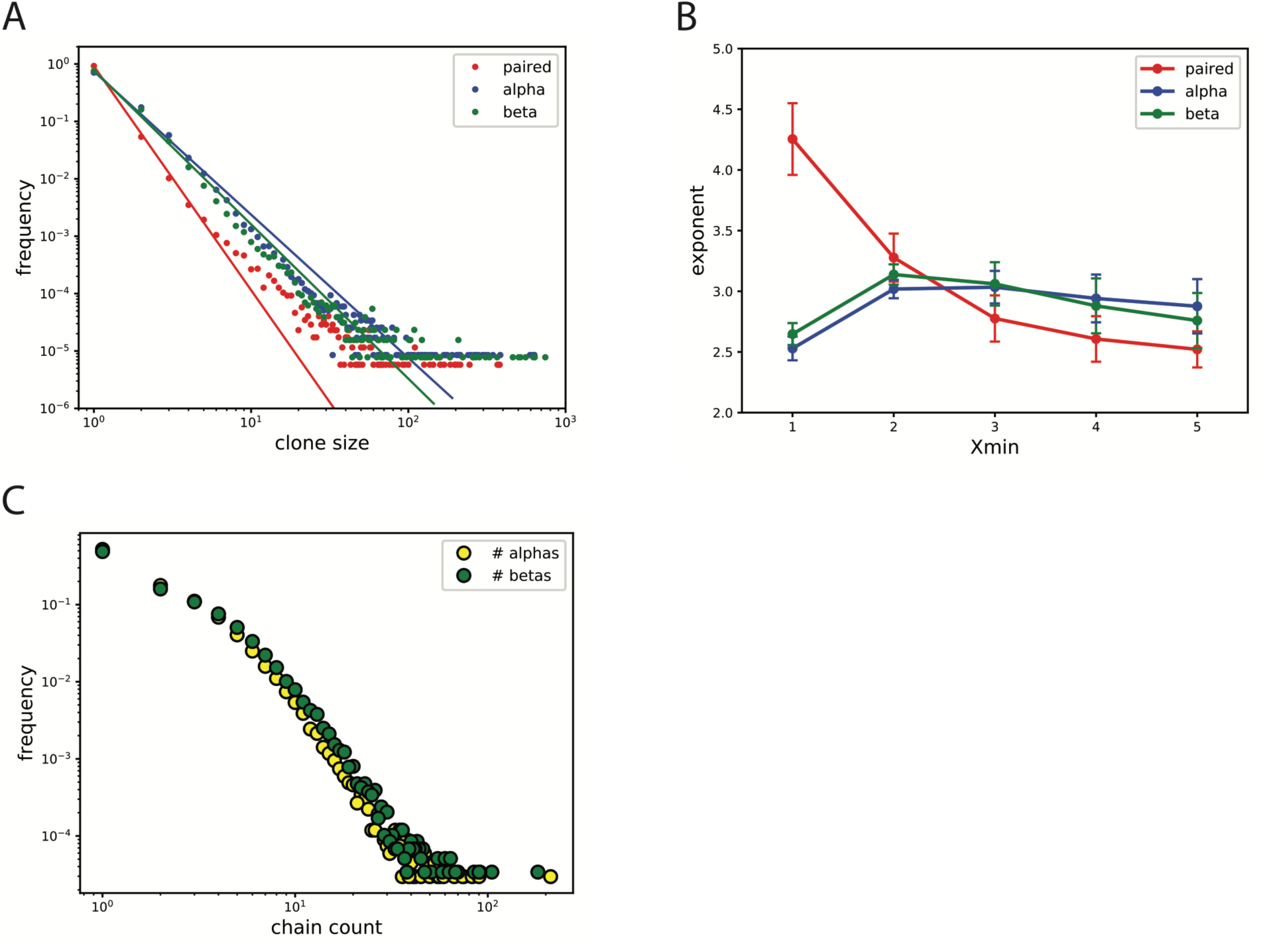
Comparison between single chain and paired αβ repertoires. A) Clone size distribution of paired αβ vs single α and β chain CDR3 sequences. We employed a maximum likelihood approach to infer the best fit of the power law distribution. Differences in repertoires suggest non-random αβ pairing. B) Exponent values for *X*_*min*_ ∈ [1,2,3,4,5], where X_min_ is a minimum clone size used in estimating the best power law fit. Change of X_min_ results in differing exponent estimates suggesting deviations from a power law behavior at small clone sizes. Significant differences in exponent between paired αβ and single chain repertoires are observed at X_min_=1 (Wilcoxon signed rank p-value=2.2×10^−2^). C) Non-unique αβ pairing. We count the number of unique α CDR3s each β chain paired with and vice versa, and looked at the probability distribution of those counts. While the majority of αβ pairings are unique, we observe low frequencies of α and β chains that pair with multiple unique β and α chains, respectively.

We quantified the differences in single vs paired αβ repertoires by fitting the power law curve and estimating the exponent (γ) using a maximum-likelihood inference approach in conjunction with Newton-Raphson numerical optimization (see Methods; Eq 8). To avoid potential biases by dominant small clone sizes, we varied the minimum clone size (*X*_*min*_) in our inference method. We report exponent estimates corresponding to *X*_*min*_ values ranging between 1 and 5, showing that (i) exponent values change with varying *X*_*min*_; (ii) paired αβ power law exponent is, on average, larger for *X*_*min*_ = 1 than that of single α and β (Wilcoxon signed rank p-value=2.2×10^−2^); and (iii) for *X*_*min*_ > 1, the differences in exponent diminishes (Figure 2B, Sup Fig. 2). These results suggest that rare clones drive the discrepancies between the three distributions. Interestingly, we observe a higher frequency of unique paired αβ clones as compared with single chain clones, providing further evidence for non-monogamous pairing of α and β chains. Additionally, we counted how many unique α chains each β chain is paired to in our dataset, and vice versa. Surprisingly, the distribution of paired counts for both chains exhibited a similar power law distribution (Fig. 2C; Sup Fig. 3). While the majority of chains exhibit unique pairing, we observe cases in each subject where some chains pair with a multiplicity of differing chains.

### Non-random associations of gene usage across the α and β chains

We next examined, whether associations between α and β gene usage were responsible for driving the differences between paired and single chain repertoires. We quantified the associations of gene usage between α and β chains, as well as between αβ CDR3 lengths, by estimating mutual information (MI), a self-equitable measure of association between two variables^25^, using a bootstrap procedure to correct for finite-sample biases (see Methods; Eq 14)^26^. We observe a consistent pattern of non-zero MI between each αβ gene pair, with higher MI for αV- βV and αJ-βV than for αV-βJ and αJ-βJ (Table 1). In comparison, MI for αCDR3-βCDR3 lengths is consistently close to zero in all subjects, suggesting no preference towards particular CDR3 length combinations.

**TABLE 1.**
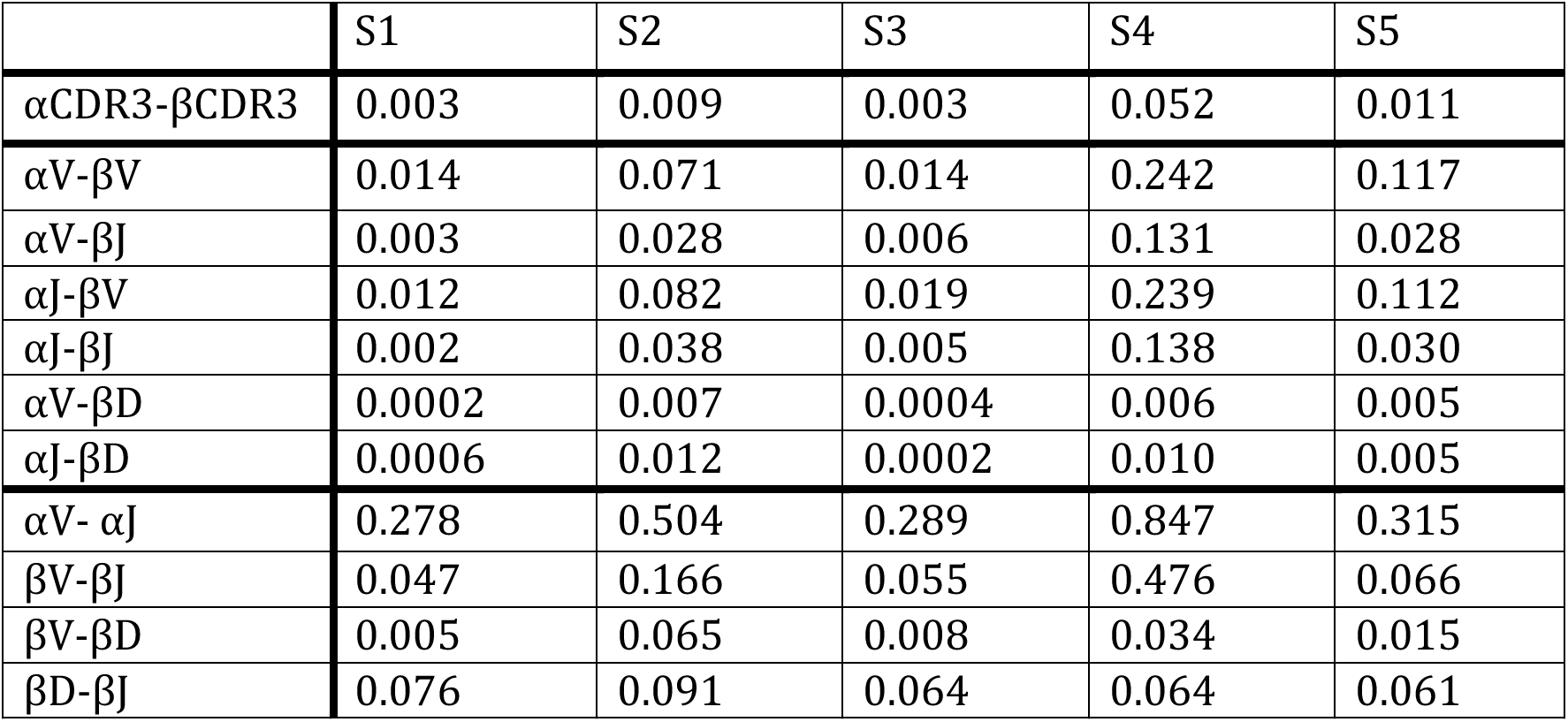
Mutual information (MI) estimates between V(D)J gene pairs within and across αβ chains, and CDR3 lengths for each subject. We applied a bootstrapping correction method to account for different sample sizes. The MI estimates reveal negligible associations between αβ CDR3 lengths, with higher MI for Subject 4, which could be driven up by clonal expansions. αβ gene pairs, particularly αVβV and αJβV, exhibit higher MI than that of CDR3 lengths suggesting that in addition to clonal expansions, there may be other unknown gene selection mechanisms during T cell maturation. In addition, MI estimates of gene pairs within the chain are higher than MI across the chains suggesting further gene selection biases during V(D)J recombination.

|  | S1 | S2 | S3 | S4 | S5 |
| --- | --- | --- | --- | --- | --- |
| αCDR3-βCDR3 | 0.003 | 0.009 | 0.003 | 0.052 | 0.011 |
| αV-βV | 0.014 | 0.071 | 0.014 | 0.242 | 0.117 |
| αV-βJ | 0.003 | 0.028 | 0.006 | 0.131 | 0.028 |
| αJ-βV | 0.012 | 0.082 | 0.019 | 0.239 | 0.112 |
| αJ-βJ | 0.002 | 0.038 | 0.005 | 0.138 | 0.030 |
| αV-βD | 0.0002 | 0.007 | 0.0004 | 0.006 | 0.005 |
| αJ-βD | 0.0006 | 0.012 | 0.0002 | 0.010 | 0.005 |
| αV- αJ | 0.278 | 0.504 | 0.289 | 0.847 | 0.315 |
| βV-βJ | 0.047 | 0.166 | 0.055 | 0.476 | 0.066 |
| βV-βD | 0.005 | 0.065 | 0.008 | 0.034 | 0.015 |
| βD-βJ | 0.076 | 0.091 | 0.064 | 0.064 | 0.061 |

Interestingly, Subject 4 exhibits higher MI for all gene pairs as well as CDR3 lengths compared to other subjects. This suggest that non-zero MI values could be, at least partially, driven by post-thymic T cell selection and proliferation due to the unusually large clones observed in Subject 4. MI for gene pairs is higher than those of CDR3 length pairs, suggesting contributions beyond clonal expansions to the observed non-random associations. We also estimated MI for gene pairs within the same chain, and found MI higher than that of gene pairs across the chains. Furthermore, associations between V and J genes of the α chain are stronger than between the genes of the β chain. This result indicates the possibility that the biases imposed on the α chain during VJ recombination is stronger than those during of VDJ rearrangement in the β chain.

Additional analyses of αβ gene pair usage frequencies reveal specific gene pairs that seem to be preferably selected during T cell maturation (Fig. 3A-F). Gene usage analyses per individual subject shows that some gene pair associations are preserved across all subjects, which could not be explained by differential post-thymic clonal expansions (Sup Fig. 4). A heat map of the αβ CDR3 length usage shows no clear associations between the chains (Fig. 3G), supporting the earlier MI calculation. We note a singular clonal expansion in Subject 4 individual CDR3 length usage heat map driving the MI up (Sup Fig. 5). Furthermore, when comparing each gene usage frequency across subjects, certain genes of both chains tend to be consistently used more often (Sup Fig. 6A-D), similar to what has been reported in previous single chain studies^27–29^.

**FIGURE 3.**
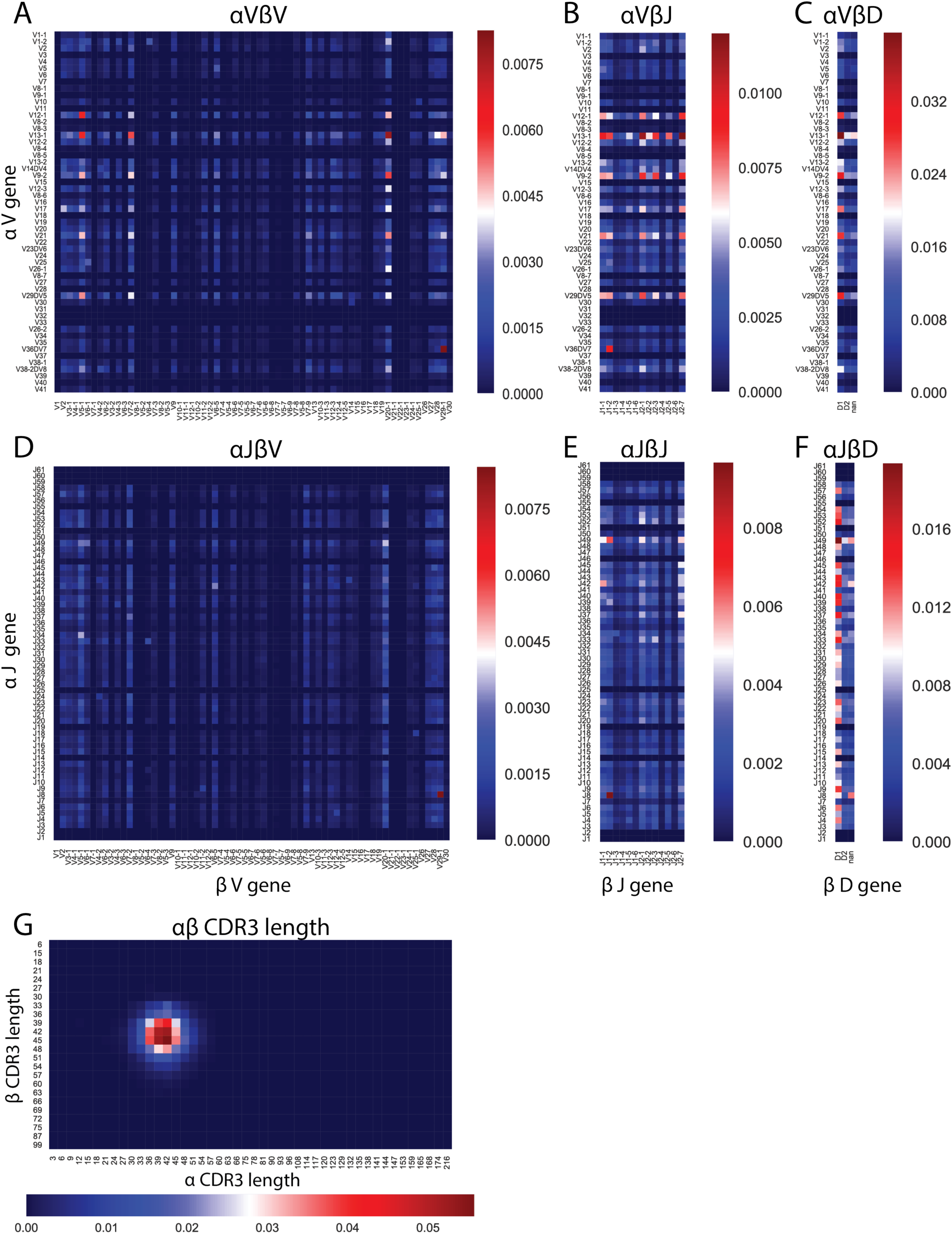
Gene usage across the αβ chains exhibits non-random associations. A-F) αβ gene usage heat maps reveal specific gene pairs used at higher frequencies, which are largely responsible for the MI values found in Table 1. Genes are listed in germ line order. G) αβ CDR3 length usage heat map reveals no preference for specific CDR3 length pairs. Note that the color scale varies between each plot.

### CD4^+^ and CD8^+^ T cell repertoires result in distinct clone size distributions

To further characterize and compare the CD4^+^ and CD8^+^ T cell repertoires across all subjects, we quantified the clone size distributions of paired αβ CDR3 sequences. We report that each subject clone size frequency distribution closely follows a power law (Fig. 4A,B) consistent with single-chain clone size distributions reported here (Sup Fig. 7) and previously^30–32^. We observe consistently differing distributions between CD4^+^ (Fig. 4A) and CD8^+^ T cell repertoires (Fig. 4B). The CD8^+^ T cell repertoire deviates from the power law behavior at the tail (Sup Fig. 8). We quantified the differences in CD4^+^ vs CD8^+^ behavior by again fitting the power law curve and estimating the exponent γ. We report exponent values for *X*_*min*_ ranging between 1 and 5 as described above. Although observed CD4^+^/CD8^+^ differences vary between individuals (Sup Fig. 9), we show that the CD8^+^ T cell exponent is, on average, smaller than that of CD4^+^ T cells (Figure 4C; Wilcoxon signed rank p-value=2.2×10^−4^). This indicates the presence of a heavier tail in the CD8^+^ as compared to the CD4^+^ distribution. Biologically, this could arise from a higher proliferative rate during clonal expansion of CD8^+^ cytotoxic T cells in an acute immune response to a given antigen, as compared with proliferative rate of the corresponding CD4^+^ T-cell population.

**FIGURE 4.**
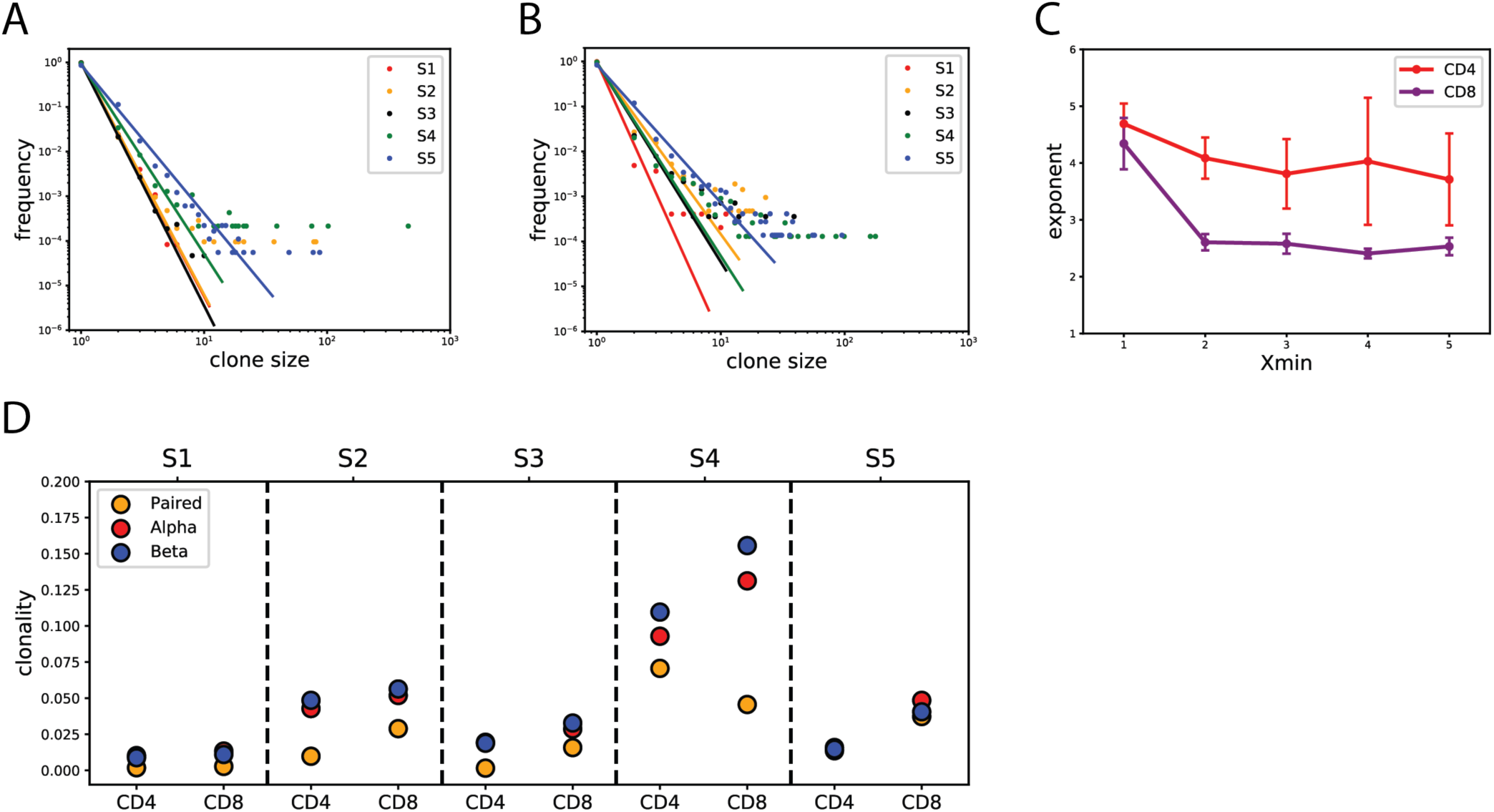
Characterization and quantification of the T cell repertoire and clonality. A) CD4^+^ and B) CD8^+^ T cell repertoires follow a power law clone size distribution characterized by linear behavior on a bilogarithmic scale plot. The power law distribution is consistent among subjects with varying exponents. The CD8^+^ T cell repertoire exhibits higher frequency of large clones compared to the CD4^+^ T cell repertoire. Maximum likelihood estimates of the power law are displayed for each subject. C) Estimated exponent values of CD4^+^ vs. CD8^+^ T cell clone size distributions for *X*_*min*_ ∈ 1,2,3,4,5. CD4 exponents are significantly higher than CD8 (Wilcoxon signed rank p-value=2.2×10^−4^). D) T cell repertoire clonality estimate comparison between paired αβ and single α and β chains for each subject. Estimates for clonality are higher in single chain data compared to paired.

### Clonality estimates differ between paired αβ, and single α and β repertoires

Next we estimated clonal abundance of paired αβ and single chain CDR3 sequences in all subjects. We calculated the clonal abundance score by subtracting normalized Shannon entropy from 1 (see Methods; Eq 12). Thus, the higher the clonal abundance score, the more homogeneous and less diverse the sample is. Our samples exhibit overall low homogeneity, with slightly higher scores in CD8^+^ compared to CD4^+^ datasets in all but one subject (Figure 4D). In addition, we compared single chain clonality scores to that of paired by splitting the paired αβ dataset into two separate, single chain datasets. We find that single chain clonality scores tend to be higher than paired, consistent with our previous observations of distinct clone size distributions of paired and single chain datasets. Subject 4 in particular exhibits an interesting behavior where the paired αβ CD4^+^ T cell repertoire clonality score is higher than that of CD8^+^. However, when considering only the single chain for this subject, we observe the opposite behavior with CD8^+^ clonality higher than that of the CD4^+^ population. This finding highlights yet another example whereby features of the paired αβ TCR repertoire cannot necessarily be inferred from single chain analyses alone.

### Existence of shared sequences across individuals

Given the finite, albeit large, number of possible human αβ pairs that could be generated by random thymic editing there may be a number of TCR sequences that are shared across individuals. The total number of T cells present in the human body at any given time is approximately 10^11^ ^3,33^, and an estimate of the number of mature T cells released into the periphery by the thymus over a lifetime (80 years) is 5×10^12^ ^34^. Given that this estimate constitutes only ~3% of all double-positive thymocytes produced in the thymus^35^, there are a total of 10^14^ productive V(D)J recombination events over a human lifetime. Experimental detection of shared sequences is also compounded by the fact that only a tiny fraction (10^−8^) of the total T cell pool is sampled. We deduce that the probability that any two sampled individuals in our study share any identical TCR sequence is approximately 10^−7^, given the number of unique TCRs observed in each subject (see Methods, Eq 10). This estimated probability suggests that we should not observe any shared sequences between individuals in our dataset. Surprisingly, we found a total of 26 shared paired αβ sequences between any two subjects and sharing was observed across most pairs of subjects. In addition, after splitting the paired αβ dataset into two separate, single chain datasets, instead of expected shared 26 single α and β sequences, we identified over 3,600 shared α and over 200 shared β chain sequences between any two individuals. Furthermore, we see a small fraction of α chain sequences shared amongst all 5 subjects, while no shared pairs are observed amongst all 5 subjects (Figure 5A, Sup Fig. 10C). After a closer look at the 26 shared paired αβ sequences we found that all 5 subjects share at least one TCR with another subject (Figure 5B, Sup Fig. 10D). This suggests that observation of shared αβ pairs is a truly biological phenomenon as opposed to a technical artifact, such as emulsion or sequencing index contamination.

**FIGURE 5.**
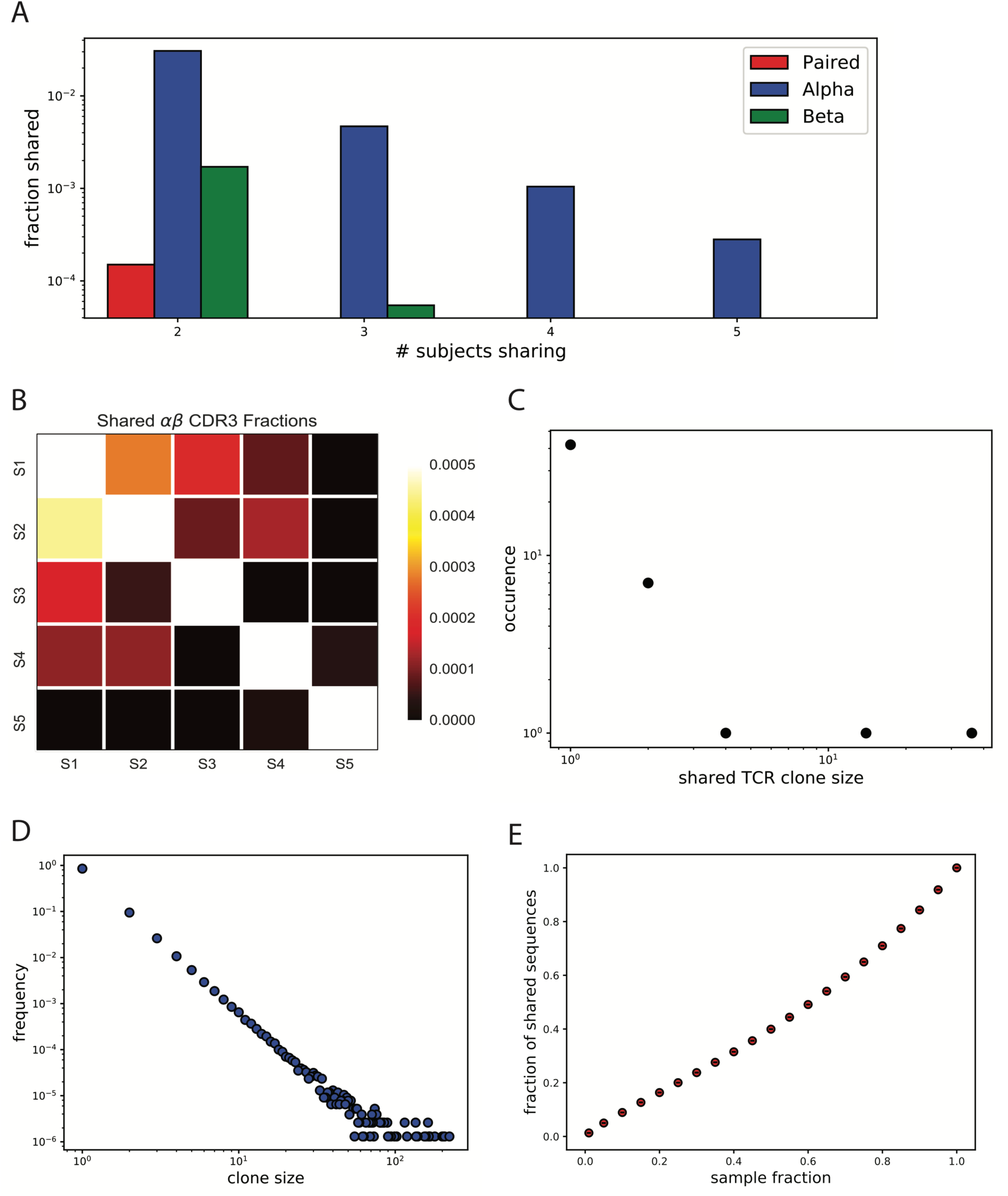
Shared sequences. A) Fractions of shared paired αβ CDR3, α chain CDR3 and β chain CDR3 sequences among subjects. B) Heat map of shared paired αβ CDR3 fractions by each subject pair. Specifically, the heat map represents a fraction of αβ CDR3 sequences of subject on the x-axis present in the subject on the y-axis. C) Clone size distribution of the 26 shared αβ CDR3s in each subject they are found. D) Simulated power law distribution (n=10^6^) resembling the T cell repertoire observed from the experimental data. Simulation was performed by creating a probability mass function (pmf) where exponent γ=3, and maximum clone size=700, and than randomly sampling from pmf generating a noisy power law distribution. E) Fractions of shared sequences observed between two equal sized sub-samples randomly sampled from the generated power law distribution (D). Rapid decrease in the number of shared sequences is observed when down-sampling.

The presence of these shared TCR sequences strongly suggests that V(D)J recombination and αβ pairing are not truly random processes, but rather that there exist strong biases in the recombination machinery that are ultimately responsible for the generation of these public clones. Furthermore, as estimates for TCR diversity rely on the assumption of random αβ pairing, these results indicate that the actual TCR repertoire may be less diverse than previously estimated. From our data, we estimated the *effective* number of human TCRs to be ~10^10^, several orders of magnitude lower than the commonly used diversity estimate of 10^15^ (see Methods).

Earlier studies reported existing biases in gene usage across individuals at a single chain level^3^, which we also observe in our dataset (Sup Fig. 6). The existence of shared paired αβ sequences suggests that biases prevail at the paired level as well. That is, certain αβ gene pairs are observed more often then others, contributing to the reduction of TCR diversity and increasing the probability of producing identical sequences in different individuals. It has been suggested that public clones should be abundant within a single individual, that is, specific TCRs that are copiously produced in any one individual are more likely to be shared across individuals^36^. To test this claim, we looked into the clone sizes of all the shared sequences and found that there is not an over representation of large clone sizes. In fact, most of the shared sequences are of clone size 1 (Figure 5C), and the distribution follows a power law, albeit with a sparse number of clone sizes. This also eliminates the possibility that observed shared αβ pairs are due to emulsion contamination by largely expanded clones.

Furthermore, to understand the probability of detecting identical sequences in the *same* individual, we performed a simulation experiment in which we generated one million sequences with the clone size distribution following the same empirical power law we obtained from the experimental data (Fig. 5D). We then randomly subsampled fractions from the distribution twice and calculated the number of sequences shared between the two subsamples. The simulation shows that even when subsampling from an *identical* distribution twice, the number of shared sequences that could be found decreases rapidly (Fig. 5E). This agrees with our experimental data in that we observe very low fractions of shared sequences between the two experimental replicates of each subject (Sup Fig. 10A, B).

## DISCUSSION

To our knowledge, we present here the largest reported dataset of paired αβ TCR sequences to date, encompassing more than 200,000 high-confidence paired receptors drawn from the peripheral blood of five healthy individuals. A critical finding of this report is that the paired αβ TCR repertoire cannot necessarily be directly inferred by observing the repertoire of one chain alone. That is, while single-chain bulk sequencing provides valuable information regarding the TCR repertoire, it is not equivalent to the paired repertoire and does not fully capture TCR diversity. The importance of αβ pairing is underscored by previous structural studies showing that receptor-antigen binding is the result of α and β peptide chain working in concert. For example, substituting a few amino acids in the antigen binding site of the α chain could either completely abolish^37^ or strengthen^38^ antigen binding. Furthermore, TCRs with identical β but different α chains can have altered antigen-MHC binding modes^39^.

We demonstrated that that T cell clone size distribution of every subject exhibits a power law behavior with heavier tails of CD8^+^ T cell distribution compared to CD4^+^, likely the result of higher proliferative activity of CD8^+^ compared to CD4^+^ T cells. While this has been previously observed at a single chain level^30–32^, we believe it to be the first confirmation of this finding with paired αβ TCRs in a high throughput fashion.

We next assessed the association between paired α and β chains, finding strong evidence for non-random and non-monogamous pairings. More specifically, non-zero mutual information between the two chains suggests that there may be substantial biases in gene usage during T cell maturation as well as post-thymic selection. We speculate, that early associations between the α and β chains likely emerge due to structural hindering during protein folding and selection processes in the thymus. We also observe associations between genes used within the same chain. It has been shown in mice that germline βV genes contain distinct promoter sequences resulting in varying levels of transcription, which could bias βV choice during recombination^40^. Furthermore, certain enhancer elements, although in the TCRδ system, were found, that regulate chromatin accessibility during V(D)J recombination influencing gene selection.

Most unexpected finding we report here is the observed 26 shared paired αβ CDR3 sequences between any two individuals. Given the estimated theoretical diversity of 10^15^ unique TCRs, and given the small sample size that could be obtained experimentally compared to the full population of 10^11^ T cells in the human body, we would expect to observe no TCR sharing among individuals. Note that V(D)J recombination occurs prior to any thymic or post-thymus selection and assuming the randomness of this process the existence of shared sequences is possible only by chance. Therefore, since the probability of TCR sharing between individuals in our dataset is 10^−7^, the existence of 26 shared paired αβ CDR3 suggests that (i) the total diversity of TCRs is smaller than 10^15^; (ii) V(D)J recombination is not a random process and unknown mechanisms must influence gene selection as well as insertion and deletion content during T cell maturation in the thymus.

Finally, to address clinical implications of our study, one may question whether differences in paired vs. single chain repertoires appear only due to rare clones. If the T cell sample is collected in a targeted way, for example from the tumor tissue, and thus only expanded clones are captured, sequencing only the β CDR3 could be enough to acquire accurate clonality estimates and changes. We argue, however, that since we find some β chains that pair with multiple α chains and vice versa, single chain measurements limits our ability to accurately quantify T cell clonality and changes. The differences in clonality measure when comparing paired vs single chain repertoire is of importance in recent cancer immunotherapy studies where the effectiveness of the treatment can be determined by tracking T cell clonal changes ^41–44^. Further studies, looking into paired αβ T cell repertoire from tumor tissues are essential for making sure that accurate clonality measures are acquired in such cases. Moreover, our public clone analysis directly shows that one can find hundreds of shared β and thousands of shared α chains among individuals with only a small fraction of those truly representing shared αβ TCRs.

In conclusion, our results stress the importance of sequencing αβ pairs to accurately describe and understand T cell diversity by emphasizing the discrepancies between single and paired αβ chain repertoires. Not only do these findings have important implications for understanding the basic biology that underlie the generation and maintenance of the TCR repertoire, but also could be particularly critical for developing successful immunotherapeutic approaches and correctly assessing patient response to treatment.

## ACKNOWLEDGMENTS

KG was funded by the Ferish-Gerry fellowship from the Watson School of Biological Sciences. JAC was partially supported by NIHGM MSTP Training award T32-GM008444 and a LIBH/NIH REACH award. GA was funded by the Simons Foundation, Stand Up To Cancer-Breast Cancer Research Foundation Convergence Team Translational Cancer Research Grant, Grant Number SU2C-BCRF 2015-001.

## METHODS

### Data Collection

PBMC samples from 5 healthy donors (three males and two females, ages 33-69) were used in our analysis, and were collected, processed and sequenced as described in Briggs *et al*, 2017 Bioarxiv^1^ (see Sup Table 1). Briefly, approximately 3.2×10^5^ viable T cells were isolated from each subject PBMC sample (Pan T Cell Isolation Kit, Miltenyi Biotec) and their T cell receptor α and β cDNA was captured in emulsion at a single cell level. For Subject 5, approximately 9.1×10^5^ viable cells were isolated and encapsulated in emulsion (EasySep HLA Total Lymphocyte Enrichment Kit, Stemcell Technologies), of which approximately 75% (6.8×10^5^) were CD3^+^ T cells. Note that Subject 5 dataset was collected at a different time, separately from Subjects 1-4. To optimize the number of droplets with 1 cell, an encapsulation Poisson rate of 0.1 cells/droplet was used. This resulted in encapsulation of approximately 1.9×10^5^ single T cells. Specific reverse transcription primers were used to target T cell receptor α and β chains along with the addition of unique molecular identifiers. The resulting cDNAs were then subjected to emulsion PCR, during which they anneal to amplified droplet barcode strands resulting in a DNA product containing droplet barcode, molecular identifier and a target sequence. The rate of droplet barcoding was tuned to a Poisson rate of 1 barcode template/droplet. After emulsion RT and PCR, the emulsion was broken and the products were purified and subjected to target enrichment PCR, where primers specific to constant regions of T cell receptors and to the droplet barcode constant sequence were used. Sequencing libraries from each emulsion were uniquely tagged with Illumina index sequences for multiplexed next-generation sequencing. The full-length TCR variable cDNA was then sequenced on Illumina MiSeq (paired-end 325+300 bp). The final raw sequence FastQ files contained cDNA sequences with their corresponding droplet and molecular barcodes, the counts of observations of each sequence and whether the sequence is an α or a β chain. In addition, we received a separate file of each droplet barcode and whether it corresponds to CD4 or CD8 T cell type determined by DNA barcode-conjugated antibody staining. Two replicate emulsions were run for each subject.

A Poisson rate of 1 barcode per droplet results in approximately 40% of cell-containing droplets containing more than one unique droplet barcode. This will modify clone size counts for small clone sizes and will have negligent effects on clone sizes >3. However, this caveat does not affect our conclusions of any analyses described here. When comparing paired vs single chain repertoires, both come from the same dataset. Thus, clone size inaccuracies will be identical and will have no impact on the comparisons. When comparing CD4 vs. CD8, the differences between the two are driven by large clones that are unaffected by the caveat. The remaining analyses do not take clone sizes into account at all.

### Data Processing

The raw sequencing data from two experimental replicates per subject were pooled together before data processing and all the sequences with occurrence of one (CONSCOUNT=1) were removed. Then, read alignment of TCR α and β sequences was performed using *Mixcr 2.2.1*^2^. Default settings were used during alignment and the following information was acquired: best V, (D), J hits, nucleotide sequences and lengths, and amino acid sequences and lengths of CDR1, CDR2, CDR3, FR1, FR2, FR3 and FR4 regions. If alignment failed for any of the sequences, those sequences were dropped by the software and did not appear in the output file. In addition, only productive sequences, those that contained a stop codon only in the FR4 region, were kept for subsequent analyses.

Our aim was to have a final dataset where one droplet barcode was associated with one sequence. However, in the raw FastQ files, some droplet barcodes appeared more than once. The next steps of data processing involved identifying those droplets and collapsing them into one. Sometimes, the repeating droplets were associated with identical sequence. In those cases such rows were collapsed into one, and occurrences of the sequences were added. In a few cases, the same droplet barcode corresponded to more than one TCR sequence. However, some of those sequences differed only by one nucleotide, which could be due to sequencing error. We performed clonality analysis with Mixcr to find those clonotypes that differ by only one nucleotide. The resulting identical clonotypes were collapsed into one as if they were identical sequences and their counts were added. The correct sequence was considered the one with the highest read count. However, some droplet barcodes that were associated with more than one sequence still remained and were thus removed from the dataset. This could be due to multiple cells per droplet or allelic inclusion. The remaining one droplet-one sequence samples were further processed by finding αβ pairs via the barcode droplet and finally, the dataset were split by CD4/CD8 type (Sup Table 1).

### Power Law curve fitting

Inferring power law exponents from empirical data is known to be non-trivial due to severe biases incurred by linear regression on bilogarithmic scales^3^. To fit the power law distributions and to accurately infer the power law exponent we employed a maximum likelihood framework with an iterative numerical optimization method (Newton-Raphson).

In general, the power law probability distribution of clone sizes with exponent *γ* is formally described by equation 1,

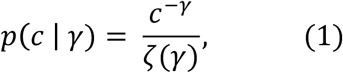

where *c* is the clone size, and

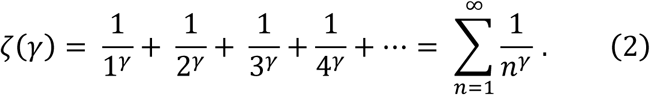

Equation 2 is the Reimann zeta function that ensures that the probability distribution is correctly normalized, i.e. 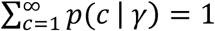. Equation 1 was further generalized by including a choice of x_min_ and x_max_ values, specifying a minimum and maximum clone size, respectively (equations 3 and 4). In our analyses, x_max_ is always set to ∞, while x_min_ is varied from 1 to 5 to show changes in the inferred exponents given the x_min_ for each condition, and observe changes in the behavior of the power law curves. Note that Equation 4 is equivalent to the Reimann zeta function when x_min_=1 and x_max_=∞,

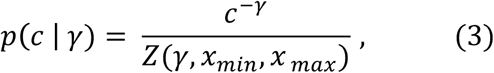

where

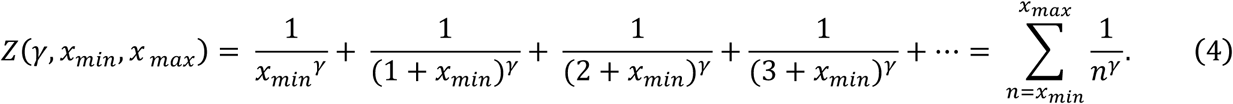

Once the data is selected within the range of x_min_ and x_max_ it consists of N numerical values {x_1_, x_2_, x_3_, …, x_N_} where xi is the clone size of TCR sequence *i*. We derive the likelihood *l* of the relevant dataset {x_1_, x_2_, x_3_, …, x_N_},

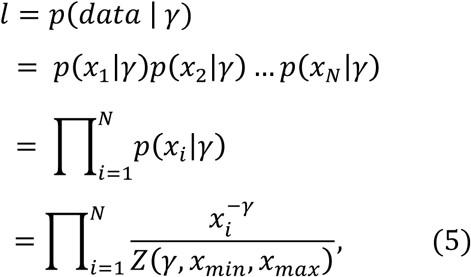

followed by the log-likelihood *L*, which is computationally more manageable

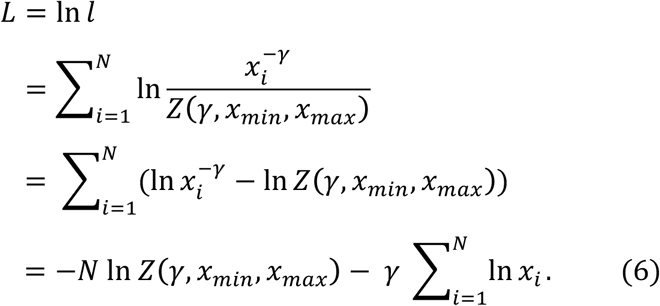

To infer the exponent γ we maximize *L*, i.e. 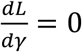. Carrying out this derivative gives us the following maximum likelihood equation where Z’ denotes differentiation of Z with respect to *γ*,

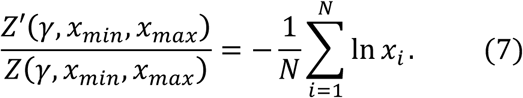

To obtain the value of exponent *γ* we solve *F*(γ) = 0, where

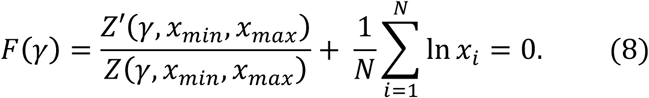

This transcendental equation is impossible to solve analytically, therefore we use an iterative numerical optimization procedure (Newton-Raphson) to solve for *γ*.

### Non-unique αβ pairing

Quality control measures were taken to ensure that observed non-unique αβ pairing reflects a biological phenomenon rather than technical artifacts. To eliminate the possibility of observing such pairs due to sequencing errors, we treated CDR3 sequences with Hamming distance 1 as identical. To reduce the possibility of an artifact due to contamination from other droplets by RNA coming from a hugely expanded clone, we removed the non-unique chain count 1 from the plots in Figure 2C and Supplementary Figure 3.

### Shared sequences

We express *f,* the probability of observing no shared TCRs between two individuals A and B, given the number of unique TCRs seen in each subject, as

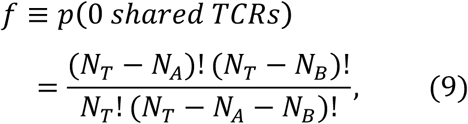

where N_T_ is the theoretical TCR diversity, and N_A_ and N_B_ are total number of unique TCRs observed in individuals A and B respectively. Given the following approximations, (i) *lnN! ≈ NlnN - N*, when *N* is large and (ii) *ln(1 - x)* ≈ *x*, when *x* is small, it can be shown that equation 9 simplifies and the probability of observing any shared TCR between two individuals is

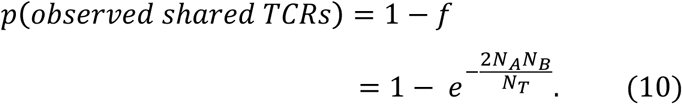

In our subjects, we observe on average ~40,000 unique TCRs per individual, and assuming that N_T_~10^15^, we find *p*(*observed shared TCRs) ≈ 10^−7^*. From equation (10) we can also estimate the effective theoretical diversity N_T_ directly from our data.

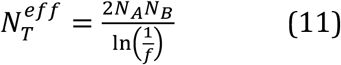

Given that subject 5, sequenced in a different batch from the rest, only shared TCR sequences with ^one other subject, we estimate the effective N_T_ to be ~10^10^, several orders of magnitude lower^ than the commonly used estimate of 10^15^.

### Clonality

Clonality measure is a clonal abundance score used to quantify how homogeneous the sampled T cell population is. The score of 1 means that every T cell in the population has an identical TCR, while a score of 0 means that every T cell has a different TCR. Higher scores are normally observed in cancer patients that have an active immune response in TILs^4^. Clonality score is calculated by subtracting normalized Shannon entropy from 1,

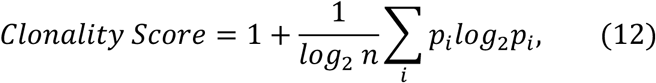

where *n* is sample size and *p*_*i*_ is a frequency of clone *i*. The clonality score is normalized to take into account different sample sizes in each subject.

### Mutual Information

To infer associations of gene usage between the alpha and the beta chains we estimated mutual information (MI) ^5^,

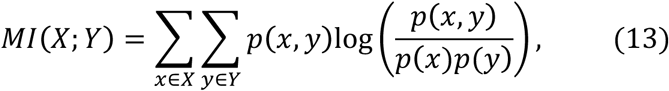

where X and Y denote the gene variables in the α and β chain.

However, limited sample sizes could bias MI estimation towards non-zero values. The true MI could only be estimated when sample size is infinity, and corrections to the frequentist-based estimate of MI scale as powers of 1/N where N is the sample size. Therefore, we perform the correction that allows us to better estimate the true MI^6^,

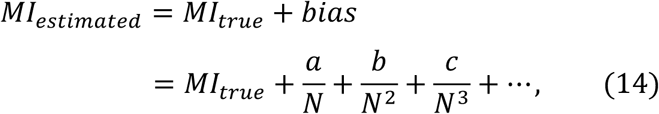

where a, b, c are constants. To estimate the true MI, we carried out a bootstrap approach: we randomly selected a fraction of sequences from the whole dataset (*fraction* ∈ [0.1, 0.15, 0.2, …, 1] and calculated MI_estimated_ for each new sample size N. MI_true_ was determined by fitting the second order polynomial curve from the function 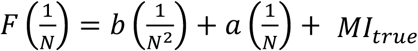 and estimating *F*(0).

### Code

All data analysis code is available as iPython Notebook at: <u>https://github.com/AtwalLab/TCRrepertoire-Manuscript</u>

All data quality control and analysis code is available in Python/bash scripts at: <u>https://github.com/AtwalLab/TCRrepertoire-Manuscript/tree/Scripts</u>

